# All-or-None Evaluation of Prediction Certainty in Autism

**DOI:** 10.1101/2022.11.17.516919

**Authors:** Seydanur Reisli, Michael J. Crosse, Sophie Molholm

**Author notes:** Corresponding author: Correspondence to Dr. Sophie Molholm.

## Abstract

The brain generates predictions to prepare for upcoming events. As life is not always 100% predictable, it also estimates a level of certainty for these predictions. Given that autistic individuals resist even small changes in everyday life, we hypothesized impaired tuning of prediction certainty in autism. To study this, EEG was recorded from adolescents and young adults with autism while they performed a probabilistic prediction task in which cue validity was parametrically manipulated. A fully predictable condition (100% cue validity) was contrasted with less predictable conditions (84, 67 and 33% cue validity). Well characterized brain potentials were examined to assess the influence of cue validity on target anticipation (contingent negative variation; CNV), the evaluation of target statistics (P3), and prediction model updating (slow wave; SW). As expected, cue validity systematically influenced the amplitudes of the CNV, P3 and SW in controls. In contrast, cue-validity effects on CNV and SW were substantially reduced in autism. This suggests that although target statistics are accurately registered in autism, as indicated by intact modulation of the P3, they are not effectively applied to generate expectations for upcoming input or model updating. Contrasting the fully predictable with the less predictable conditions, our data suggest that autistic individuals adopted an all-or-none evaluation of certainty of their environment, rather than adjusting certainty of predictions to different levels of environmental statistics. Social responsiveness scores were associated with flexibility in representing prediction certainty, suggesting that impaired representation and updating of prediction certainty may contribute to social difficulties in autism.

**SIGNIFICANCE STATEMENT:** The ability to make predictions is integral to everyday life. Yet, as life is not always 100% predictable and it is also essential to adjust the certainty of these predictions based on the current context. This study reveals that individuals with autism are less efficient in adjusting the certainty of their predictions to the level of predictability of events. Instead, they may adopt an all-or-none evaluation of certainty. Our findings reveal novel insights into the processes underlying impaired predictive processing in autism, which may open the door to developing targeted behavioral interventions and/or non-invasive brain stimulation therapies that help autistic individuals make more accurate predictions to ease social- and rigidity-based symptoms.

## INTRODUCTION

Predicting what comes next is highly advantageous for adaptive behavior and leads to facilitated processing of information (Bar, 2007; Gregory, 1980; Hohwy, 2017). Many current theories of perception propose that the brain maintains a model of the environment that produces top-down predictions of upcoming stimuli at various hierarchical stages of processing, rather than simply acting on sensory inputs (Bar et al., 2006). These predictions are associated with high certainty for predictable environments and low for volatile environments (Friston & Kiebel, 2009). For adaptive behavior, predictions and the associated level of certainty (e.g., *precision*) must flexibly be updated based on new information.

Predictive processing accounts of autism have gained popularity (Cannon et al., 2021) as they not only provide a model within which to generate falsifiable hypotheses (Friston & Kiebel, 2009), but also explanation for a diverse range of autism symptomology including cognitive-, sensory-, and motor-related characteristics (Gomot & Wicker, 2012; Van de Cruys et al., 2014). For example, problems in social communication have been attributed to reduced ability to form generative models that can be used to predict and interpret social cues (Chambon et al., 2017; Palmer et al., 2015), and resistance to change to an overly rigid predictive model (Gomot & Wicker, 2012) such that unexpected changes cause discomfort. There is mounting support for suboptimal updating of the predictive model in autism (Coll et al., 2020; Palmer et al., 2017), including evidence of slower model updating (Sapey-Triomphe et al., 2021; Soulières et al., 2011; Vishne et al., 2021), and oversensitivity to prediction errors that leads to bigger model updates in response to errors ((Karvelis et al., 2018; Van de Cruys et al., 2014), but see (Knight et al., 2020)).

In a recent study, a smaller difference in response time between conditions where cues were more versus less predictive of a target (84% vs. 16%) was observed in autism compared to controls, which was interpreted as reduced surprise in autism upon prediction violation (Lawson et al., 2017). This and similar findings (Perrykkad et al., 2021) appear counter-intuitive with clinical observations and introspective reports that autistic individuals overreact to violations of expected outcomes. In these studies, however, conclusions are based on comparison between conditions for which the cue is never fully predictive. Arguably, if resistance to change and rigid adherence to routines results from intolerance to any violation of predictions, a 100% predictable condition provides an important baseline against which to assess the magnitude of the surprise response. However, no study that we are aware of has juxtaposed a fully predictive condition with less predictive conditions.

To better understand the representation of certainty of predictions in autism, we designed a probabilistic task where an initially fully stable environment was achieved with 100% cue validity, while three further levels of cue validity (i.e., 84%, 67% and 33%) were presented later. Using this task accompanied by EEG recordings, we tested the representation of different levels of cue validity in individuals with autism. In the control group we expected a more-or-less linear relationship between the primary dependent measures and cue validity, indicating that certainty is represented in a graded manner. In contrast, given that autistic individuals over-react to deviations from expectations (Frith, 2003; Lord et al., 2012), we expected the autism group to show bigger differences in behavioral and brain responses compared to controls between a fully predictable condition (i.e., 100% cue validity) and a slightly less predictable condition (i.e., 84% cue validity). On the other hand, we expected less clear differentiation among the less predictable conditions (e.g., across 84%, 67% and 33% cue validities), consistent with findings in the literature of reduced differential responses to changes in less versus more stable environments (Lawson et al., 2017; Perrykkad et al., 2021). Well-characterized Event Related Potentials (ERPs) allowed us to assess the evaluation of the cue-target statistics (e.g. P300), and how individuals used these statistics to modulate their expectations in preparation for upcoming targets (e.g. CNV).

## METHODS

### Experimental design and statistical analysis

#### Participants

Nineteen individuals with autism (8 left-handed, mean age: 19.6 ±2.7 years old) and 21 Intelligence Quotient (IQ)- and age-matched control subjects (all right-handed; mean age: 20.7 ±2.32 years old) participated in the study, all aged between 16 and 28 years (Table 1). Autism diagnoses were made using the Autism Diagnostic Observation Schedule, Second Edition (ADOS-2) (Lord et al., 2012), the Autism Diagnostic Interview-R (Lord et al., 1994), and expert clinical judgment by a licensed psychologist at the Human Clinical Phenotyping Core of the Rose F Kennedy Intellectual and Developmental Disability Research Center (RFK IDDRC) at the Albert Einstein College of Medicine.

**TABLE 1:**
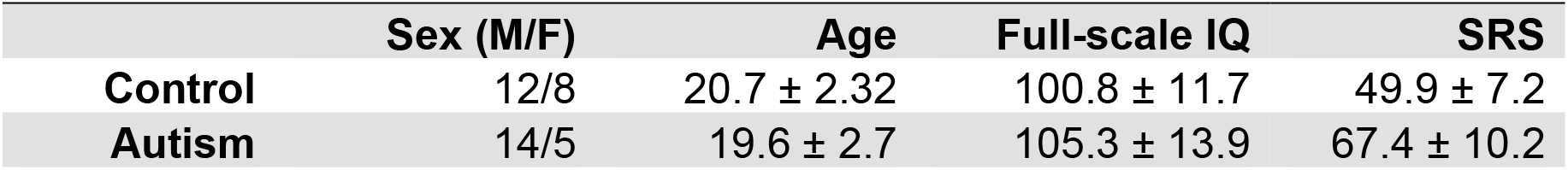
Participant Demographics. Mean and standard deviation values are reported for age, full-scale IQ, and Social Responsiveness Scale (SRS). The Full-Scale IQ was based on Wechsler Abbreviated Scale of Intelligence (WASI).

|  | Sex (M/F) | Age | Full-scale IQ | SRS |
| --- | --- | --- | --- | --- |
| Control | 12/8 | $20.7 \pm 2.32$ | $100.8 \pm 11.7$ | $49.9 \pm 7.2$ |
| Autism | 14/5 | $19.6 \pm 2.7$ | $105.3 \pm 13.9$ | $67.4 \pm 10.2$ |

Participants were recruited without regard to sex, race, or ethnicity. Exclusionary criteria for both groups included a performance IQ below 80; a history of head trauma; premature birth; a current psychiatric diagnosis; or a known genetic syndrome associated with a neurodevelopmental or neuropsychiatric condition. Attention deficit/hyperactivity disorder (ADD/ADHD) was not used as an exclusion criterion for the autism group, given its high comorbidity with autism. Exclusion criteria for the control group additionally included a history of developmental, psychiatric, or learning difficulties, and having a biological first-degree relative with an autism diagnosis. Participants who were on stimulant medications were asked to not take them at least 24 hours prior to the experiment.

#### Neuropsychological and clinical testing

IQ was measured via the Wechsler Abbreviated Scale of Intelligence (Simard et al., 2015). To quantify autism-related characteristics, both groups of participants completed the Social Responsiveness Scale-2 (SRS-2) (Constantino, 2013) which has five subscales (i.e., Social Awareness, Social Cognition, Social Communication, Social Motivation, and Restricted Interests and Repetitive Behavior (RRB)). We used the self-report SRS-2 total t-scores to assess correlations with participant EEG and Reaction Time (RT) measures.

Independent paired t-tests showed no significant group differences for age [t(44) = 0.95, p=0.34] or full-scale IQ [t(40) = −0.40, p=0.69]. Among various sub-domains of the Wechsler Intelligence test, only one domain, the processing speed index (PSI), showed a significant group difference [t(30) = 7.59, p<0.01] revealing that autism group was slower in processing information. As expected, the autism group had higher SRS-2 scores than the comparison group [t(33) = −8.48, p<0.01], as well as on each of the SRS-2 sub-domains.

#### Sequential Probabilistic Task

We designed a task to probe the ability to adjust prediction certainty based on changing probabilities in the environment.

##### Stimuli

Visual stimuli were presented to the participant, one at a time, on a computer screen at a viewing distance of 65 cm in a dimly-lit room. Stimuli consisted of basic shapes presented in gray on a black background for 100 ms, with an 850 ms interstimulus interval (ISI). Participants performed a target detection task in which they responded as quickly as possible to the final item of a target-sequence. A target-sequence was either three arrows, the first upward-facing, the second rightward-facing, and the final downward-facing, or three parallelograms, the first left-tilted, the second straight, and the final right-tilted. The stimuli in these sequences are referred to as cue1, cue2, and target (Fig. 1A). When patterns were not completed, a circle, diamond, or triangle shape was presented instead, which we refer to as an invalid item. These shapes were also used as *fillers*, represented once or twice after invalid items or targets. To ensure that participants were responding to the shape sequence and not just the final shape in the sequence, catch trials in which the final shape was presented after filler shapes were also included.

**FIGURE 1:**
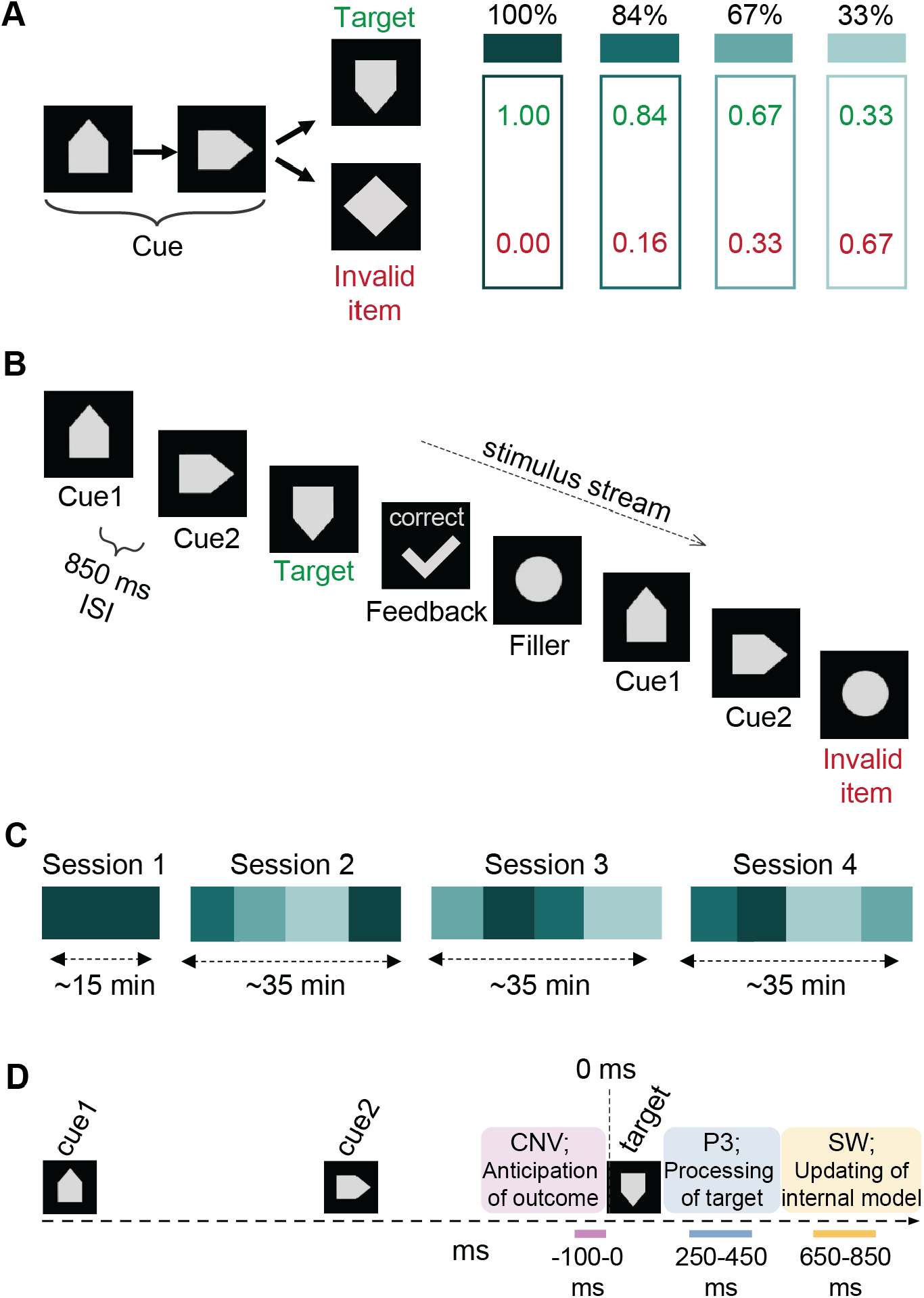
The Sequential Probabilistic Task. (A) Participants respond to target sequences of stimuli while the probability of sequence completion is manipulated at four levels. Stimuli consist of basic shapes presented sequentially to the participant. The two possible target sequences: A sequence of 3 arrow or 3 parallelogram shapes are presented in specific orders. The participant’s task is to respond after targets with a mouse click while withholding the response after invalid items (B) A sample sequence in time from the experiment is provided as an example. The subject responds with a mouse click after completion of the three pattern items, followed by a feedback message appearing on the screen. (C) The order of probability conditions throughout the experiment is shown for a sample participant. (D) Conceptual illustration of the temporal dynamics of evoked responses of interest: CNV, P3, and SW.

##### Probability conditions

Throughout the experiment, the probability that a target-sequence was completed varied across four levels, in ~10 min blocks (Fig. 1C). Pattern initiations, always represented by cue1 of the pattern followed by cue2, were completed with the target stimulus 100%, 84%, 67% or 33% of the time, comprising four cue validity conditions (Fig. 1A). Participants were not informed of the probability condition they were in or when it changed. The two target-sequences were presented with equal probability within a given probability condition.

##### Blocks

Stimuli were presented in mini blocks of ~1.5 minutes, separated by pauses during which time participants could rest. Each mini block was composed of 24 pattern initiations (cue1 followed by cue2) (see Table S1 for more). Pattern initiations were completed with the target 24, 20, 16 or 8 times depending on the probability condition. Participants pressed the mouse key to initiate the next mini block. Blocks of a given probability condition were composed of between 4 and 6 mini blocks.

##### Instructions Part 1

The following instructions were printed on the screen in four parts, both for remote familiarization and the first experimental session:

> *“You will see a shape in the middle of the screen. The shape will change about every second. Sometimes 3 consecutive shapes appear in the orders below, which we call a pattern. There are two target patterns: (pattern shapes were shown to the participant below this sentence). Your job is to touch the screen (or press the mouse button) after Pattern 1 or Pattern 2 is completed. Try to be both quick and accurate. Remember, you should respond after the pattern is completed. You can ignore any other shape. Let’s practice!”*

##### Remote Familiarization

To briefly familiarize participants with the stimuli and task prior to the experiment, we remotely presented the task (100% cue validity condition only) for six minutes using the Neurobehavioral Systems mobile app on their smart phone or tablet, one day before the experiment.

##### Experiment sessions

The experiment was composed of four sessions performed on a single day, separated by 15-30 minute breaks (Fig. 1C). In Sessions 1 and 2, the probability conditions were presented in the same order to all participants, whereas in Sessions 3 and 4, probability condition order was pseudo-randomized. Session 1 consisted of 7 mini blocks of 100% cue validity condition. In Session 2, conditions were presented in the order of 84%, 67%, 33%, and 100%. Participants usually took a lunch break after Session 2, while taking a ~15-minute break between Session 3 and Session 4. In Sessions 3 and 4, probability conditions were presented in a pseudo-randomized order (sample order is shown in Fig. 1B). The initial 100% condition, presented during remote familiarization and Session 1, was designed to establish strong cue-outcome associations. This might correspond to never broken rules that individuals with autism seek in adhering to strict routines in their everyday life.

##### Instructions Part 2

At the end of the first session, participants were informed that going forward, the cues would not always be followed by the target, and that in these cases they should withhold their response.

##### Feedback

To keep the participant on-task, visual feedback was provided: “correct” for responses to targets that fell within the response window of 100 to 950 ms; “miss” if they did not respond within 950 ms of the target; “too early” for responses occurring within 100 ms of target presentation (assumed to be anticipatory); and “wrong” for responses to a non-target. Feedback text was accompanied by an icon (a “✓” for correct, “x” for wrong, “!” for miss or too early). The feedback stimulus was presented for 200 ms.

#### EEG data collection and pre-processing

Continuous EEG was recorded from 160 scalp electrodes at a rate of 512 Hz using the BioSemi ActiveTwo system (BioSemi B.V., Amsterdam, Netherlands). Biosemi replaces the ground electrodes that are used in conventional EEG systems with two separate electrodes: Common Mode Sense (CMS) and Driven Right Leg (DRL) passive electrodes. These two electrodes create a feedback loop, thus rendering them as references. Data were down-sampled to 128 Hz for subsequent analyses, to reduce computing demands. EEG data were pre-processed using Matlab and eeglab (Delorme & Makeig, 2004) on local computers or remote cluster computing via Neuroscience Gateway (Sivagnanam et al., 2013). Data were high-pass filtered at 0.75 Hz. The 60 Hz line noise was removed using CleanLine function of eeglab, run twice with a window and step-size of four. Channels that were two standard deviations away from the average power spectrum in the 0.1-50 frequency band were rejected.

Infomax Independent Component Analysis (ICA) was used to remove potential non-brain related activity, mainly eye-movement related muscle artifacts. For each Independent Component (IC), the iclabel program (Pion-Tonachini et al., 2019) was used to calculate the probabilities for that IC belonging to the seven different IC categories including Brain, Muscle Noise, Eye Noise, Heart Noise, Line Noise, Channel Noise, and Other. A total noise metric was created via summation of muscle-, eye-, heart-, line-, channel-related noise probabilities. An IC was excluded only if it met both of the following criteria: 1) had more than 50% chance for the noise category, 2) had less than 5% chance of the brain category. This led to an average of 5 ICs being rejected among the top 20 ICs (i.e., the ICs that accounted for the majority of the signal). Three of these on average had more than 50% chance of being a component related to eye blinks or movements. The channels that were rejected prior to ICA were interpolated using linear interpolation method. After referencing data to the average of two scalp channels that are near the right and left mastoids (i.e., E17 and B18 on BioSemi 160 System). For P3 and SW analyses data were epoched between −100 and 950 ms with respect to stimulus onset, with the first 100 ms of the epoch serving as baseline. For the CNV analyses data were epoched between −100 and 950 ms with respect to the second cue, with the first 100 ms serving as baseline. Noisy trials were rejected based on a custom script that rejects trials with amplitudes that are more than three standard deviations away from the mean of maximum global field power amplitudes for each trial type. After that, trails were averaged for each stimulus type.

### Data analyses

EEG, reaction time, accuracy, and clinical data were analyzed in Matlab and Python using custom libraries and scripts. We assessed the effect of cue validity on three well-characterized Event Related Potentials (ERPs) to the temporal dynamics of predictive processing in response to changing environments: The contingent negative variation (CNV), a slow negative-going ERP that typically systematically varies in amplitude with the certainty of target expectation (Thillay et al., 2016) and represents anticipatory brain activity involved in preparing a response to a temporally predictable target (Brunia, 2003), and the P3 (aka P300), a positive-going ERP associated with target detection and evaluation that occurs in response to a target, and varies in amplitude with respect to target probability (Bidet-Caulet et al., 2012; Polich, 2007, 2012). While the P3 allowed us to assess the evaluation of the cue-target statistics, the CNV provided information about how individuals used these statistics to modulate their expectations in preparation for upcoming targets. In addition, we measured the slow wave (SW) to index updating of the internal model. Selection of the temporal windows and scalp regions used for analysis of each of these components was informed by the literature and modified if needed based on inspection of the specific timing and topography of the response of interest, without regard for experimental condition or group. The CNV was measured as the average amplitude over the 100 ms window preceding the onset of the imperative stimulus (the target or the invalid item), from a centrally placed electrode (one anterior to the classic Cz location) (Thillay et al., 2016). The P3 was measured as the average amplitude between 250-450 ms (+/-100 ms from the 350 ms peak) at Pz (Polich, 2007). The Slow Wave response (SW) was measured as the average amplitude between 650-850 ms (+/-100 ms from the 750 ms peak) following the target at electrode Fpz (de Gee et al., 2021; Sambrook et al., 2018). While measuring P3 and CNV responses was an apriori decision, the SW was a post-hoc analysis (see the Results and Discussion sections for justification). For behavioral analyses, RT, percent hits, and false alarms were calculated for each participant for each cue validity condition, and subsequently averaged per participant group. In our tasks, in line with prior work, RT was expected to be faster with increasing cue validity across conditions (Lawson et al., 2014; Thillay et al., 2016).

For statistical analyses of the single-trial relationship between cue validity and ERP amplitude, we conducted linear mixed-effects models using statsmodel package in Python (Seabold & Perktold, 2010). Models were fit using a maximum likelihood criterion defining subjects as a random factor. ERP amplitudes were the numeric dependent variable. Group was a dummy-coded fixed factor.

To test the hypothesis that flexibilty in certainty of predictions relates to social responsiveness, we conducted correlation analyses between clinical scores and our primary EEG measures. We took the difference between 84% and 33% conditions as an index of a participants’ ability to differentiate between different probability conditions (e.g., prediction flexibility index). We then performed Pearson’s correlation between this index and social responsiveness (as measured by SRS-2).

Our hypothesis-driven analyses risks oversight of potentially informative patterns in these rich high-density EEG data. Therefore, exploratory analyses were performed on the full data matrix to serve as a hypothesis generation tool for future studies. To this end, running statistical tests were carried out across all channels and time points (Molholm et al., 2002; Morie et al., 2014). We displayed the results of running t-tests between 84% and 33% conditions as an intensity plot (e.g. Fig. S1). The x and y axes represent time and electrode location respectively, while the heatmap represents the p value for each data point. Called statistical cluster plots (SCPs), this provided us a simple approach for testing for differences between a given pair of experimental conditions across the entire data matrix. Following the rationale of earlier approaches (Molholm et al., 2002; Morie et al., 2014), type 2 errors were minimized by only displaying significant differences when at least three consecutive time points (from data downsampled to 128 Hz, thus representing a 22ms time window) and three neighboring channels (significance was required for at least three out of eight surrounding channels in the Biosemi 160 system) were significant.

### Code accessibility

All code is available online under three repositories: 1) The code that was generated for stimulus presentation using the Presentation software^®^ is available at https://github.com/seydareisli/splt; 2) the Matlab code that was used to process data is available at https://github.com/seydareisli/mat; 3) the Python code that was used for visualization and figures is available at https://github.com/seydareisli/viz.

## RESULTS

We designed a sequential probabilistic task where participants responded to the completion of three sequentially presented shapes (e.g., three arrows, the first upward-facing, the second right-facing, and the final downward-facing; aka cue1, cue2 and target) while parametrically manipulating sequence completion at four levels: 100%, 84%, 67%, and 33%. The effects of probability condition and autism diagnosis on brain signals (i.e., P3 and SW after targets; CNV after cue2) and RT were examined to understand how different levels of certainty in predictions (e.g., stimulus predictability) is represented in the brains of individuals with autism.

### Electrophysiological data

To assess if brain potentials reliably modulate as a function of probability and whether this significantly differs by group, we performed three separate linear mixed effect models for P3, CNV, and SW. ERP amplitudes were best fit by a linear mixed effect model by including an interaction term between group (control and autism) and probability condition (100%, 84%, 67%, 33%). Post-hoc mixed models were conducted for each potential pairwise comparison (100-84%, 100-67%, 100-33%, 84-67%, 84-33%, 67-33%) to unpackage the significant main effects and group-by-condition interactions. Results are reported below and summarized in Table 1 and in supplementary Table 2.

CNV: In both the autism and control groups, a CNV was observed just prior to onset of the imperative stimulus (target or invalid item). The CNV, which had a central negativity, was most prominent in the 100 ms prior to stimulus onset (Fig. 2, S2). In the control group, this amplitude appeared more negative-going as cue validity decreased. In the autism group, CNV amplitude appeared highly similar across the three less predictable conditions (i.e., 84%, 67%, 33%), while these clearly segregated from the 100% condition. Statistical testing of the data revealed a significant effect of condition (ß=1.54, SE=0.18, p<0.01) and a group-by-condition interaction (ß=-0.64, SE=0.26, p=0.01), but no significant effect of group (ß=1.38, SE=10.99, p=0.90). Post-hoc follow-up tests revealed that the condition effect was driven by all pairwise comparisons except the 84-67% comparison (the two more ambiguous conditions), whereas the group-by-condition interaction showed smaller differences in the autism compared to the control group for the comparisons of the 33% to the other conditions.

**FIGURE 2:**
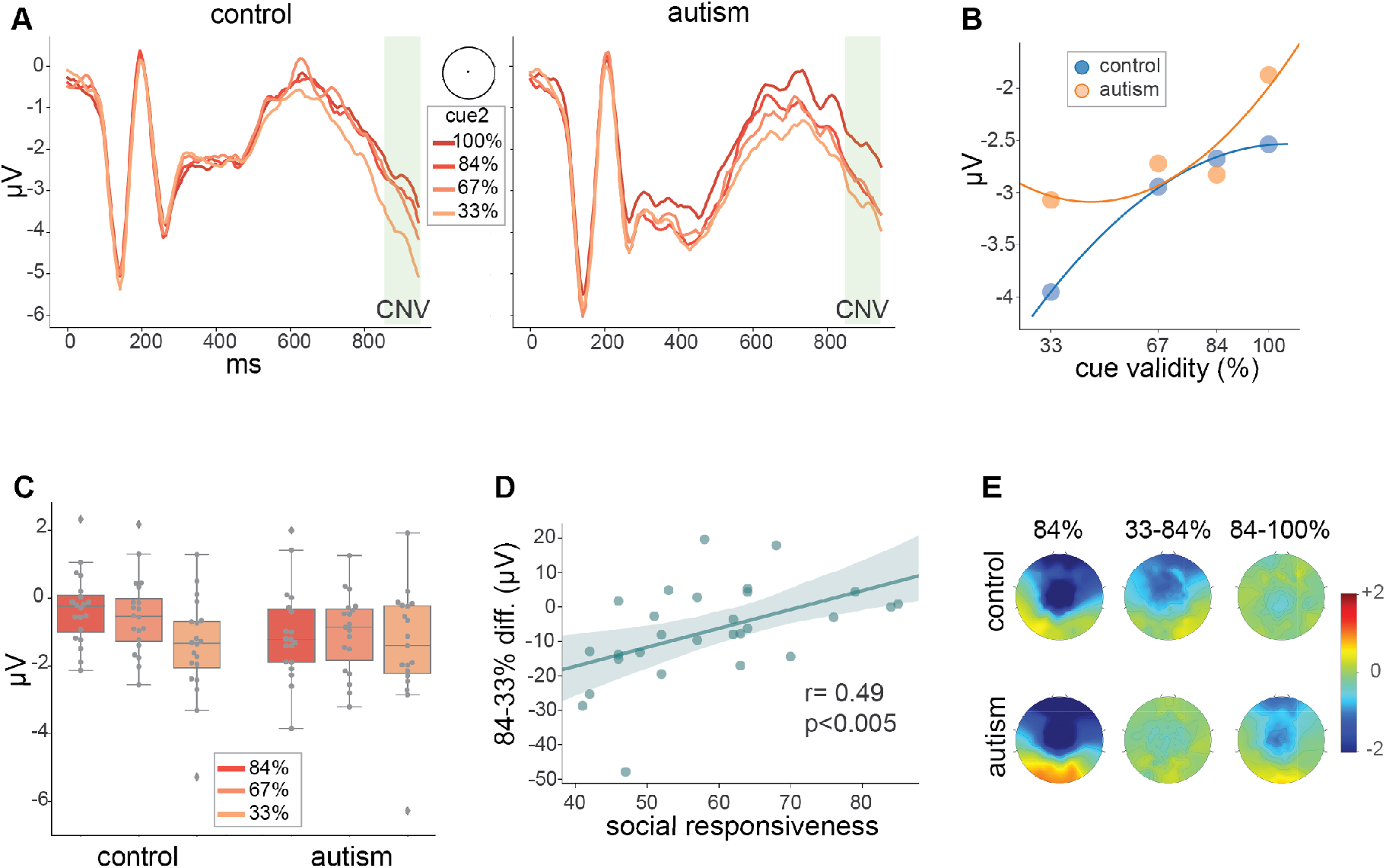
CNV. (A) ERP waveforms showing responses timelocked to cue2 at Cz for each of the cue validity conditions. The CNV time window is highlighted in green (100 ms prior to target onset). (B) Average CNV amplitude across the four validity conditions measured at Cz with second-order polynomial fits shown for each group. (C) CNV amplitudes across 84%, 67%, 33% conditions normalized to 100% condition, error bars showing 95% confidence intervals. (D) Pearson’s correlation between Social Responsiveness Scores and CNV flexibility index (difference waveform between 84% and 33% conditions). (E) CNV topographies for 84% condition (left), difference between 33% and 84% conditions (middle), and difference between 84% and 100% conditions (right).

P3: Both groups exhibited a typical P3 in response to target stimuli that was positive-going over posterior-central scalp and peaked at about 350 ms. Furthermore, in both groups, the amplitude of the P3 varied as a function of cue validity (Fig. 3, S2). The P3 statistical model revealed a significant effect of condition (ß=-3.19, SE=0.21, p<0.01), while showing no main effect of group (ß=-0.43, SE=9.02, p=0.96) or group-by-condition interaction (ß=0.14, SE=0.30, p=0.65). The main effect of condition was driven by all pairwise comparisons of cue validity conditions.

SW: The P3 was followed by a second phase of post-target activity that was positive going over the frontal scalp and was apparent in both groups. This was seen in the 650-850 ms window. For the control group, this response appeared to be larger and of longer duration in lower cue validity conditions, whereas there was no obvious systematic modulation of this response by condition in the autism group (Fig. 4) (see the second-order polynomial fits in Fig. 4B showing a linear versus curved relationship in controls versus autism groups). This response resembles the SW component, a brain response that is observed in cued tasks (de Gee et al., 2021; Loveless et al., 1987; Ruchkin et al., 1980), typically occurs in this same window after a target or invalid item, also has a positive-going frontal scalp distribution, and varies in amplitude with respect to cue validity. The statistical model revealed a significant effect of condition (ß=-1.44, SE=0.26, p<0.01) and a significant group-by-condition interaction (ß=1.72, SE=0.38, p<0.01), but no main effect of group (ß=-0.86, SE=11.29, p=0.94). The main condition effect was driven by all pairwise comparisons of probability conditions except the 67-33% contrast. The significant group-by-condition interaction was due to all pairwise comparisons except the 100-84% and 67-33%. Group-by-condition interactions reflected smaller differences in the autism compared to control group. In autism, the SW was of greater amplitude in the 84% compared to the 67% and 33% conditions, which contrasts with opposite pattern in controls (see Fig. 4 and Table S4).

**FIGURE 3:**
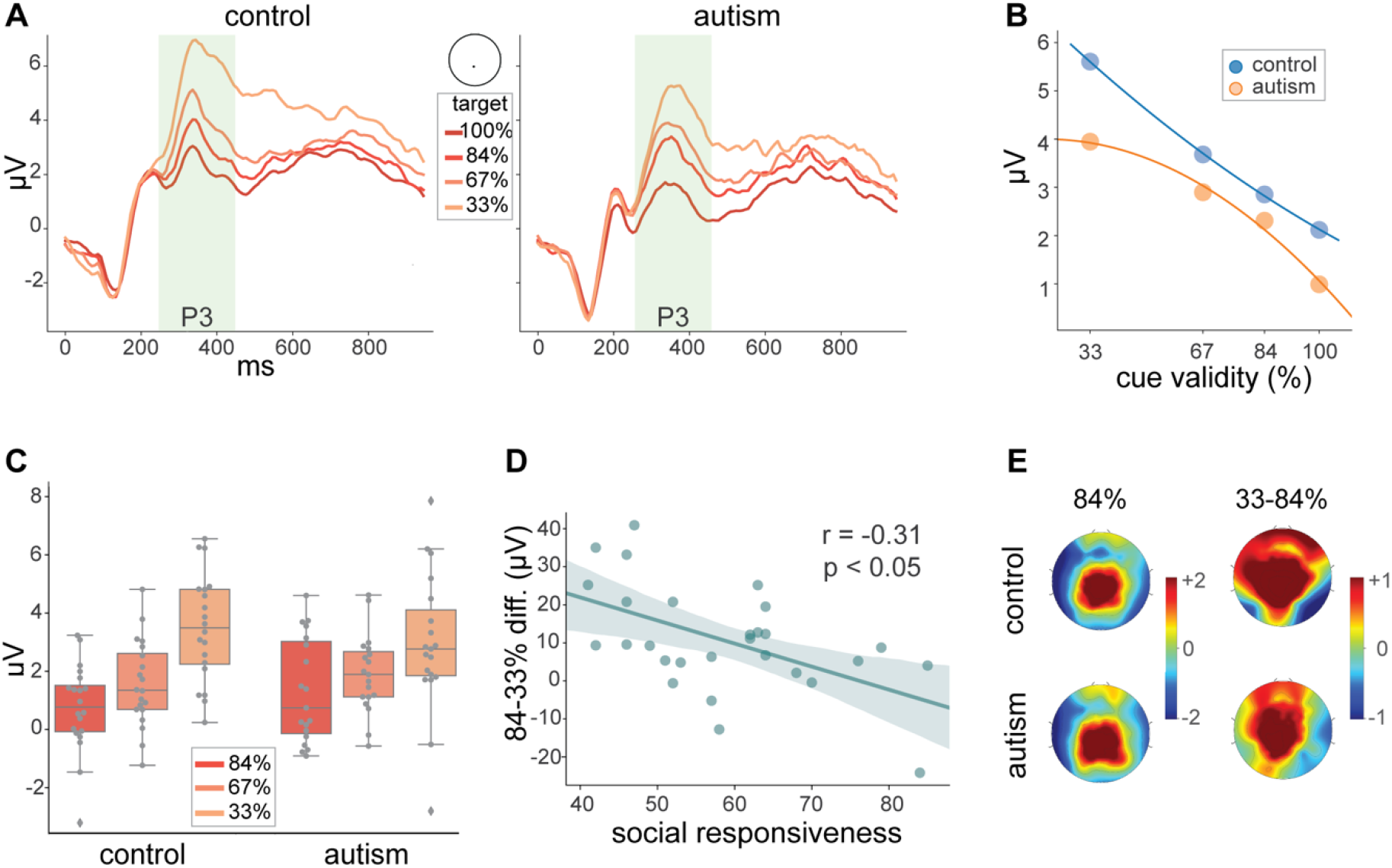
P3. (A) Target-locked ERPs at Pz; P3 time window highlighted in green panel. (B) Average P3 amplitudes for each group across the four validity conditions. Lines show second-order polynomial fits for each group. (C) P3 amplitudes across 84%, 67%, 33% conditions normalized for 100% condition, error bars showing 95% confidence intervals. (D) Correlation between Social Responsiveness Scores and P3 flexibility index (difference waveform between 84% and 33% conditions). (E) P3 topographies for the 84% condition (left) and P3 difference topographies between 84% and 33% conditions (right) are included for each group.

**FIGURE 4:**
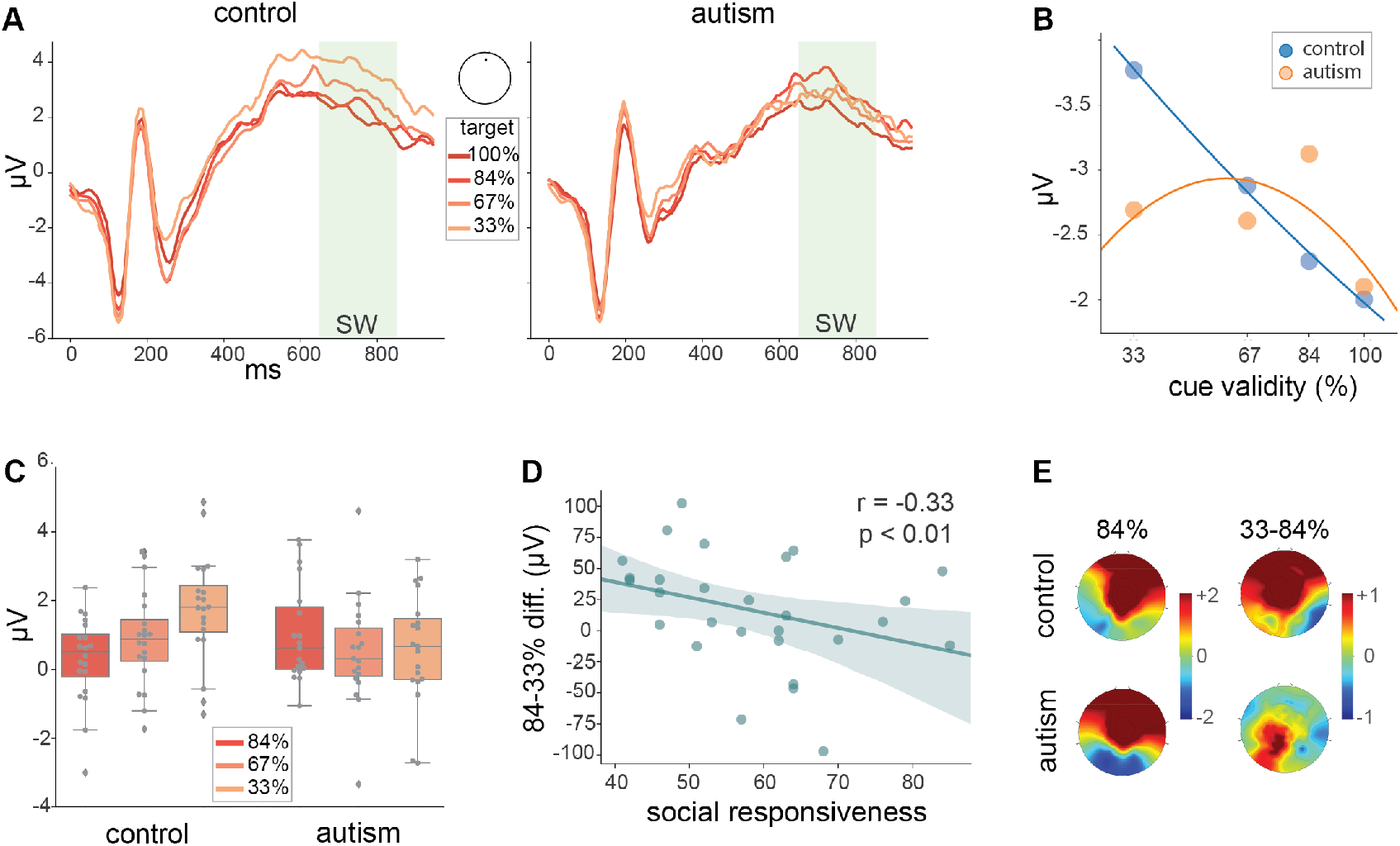
SW. (A) Target-locked ERPs at Fpz; SW time window is highlighted in green. (B) Average SW amplitude for each cue validity condition with second-order polynomial fits. (C) CNV amplitudes across 84%, 67%, 33% conditions normalized to 100% condition, error bars showing 95% confidence intervals. (D) Correlations between SRS and SW flexibility index (difference waveform between 84% and 33% conditions). (E) Topographies are included for each group for the 84% condition (left) and SW difference topographies between 84% and 33% conditions (right) are included for each group.

Statistical Cluster Plots: The SCPs contrasting the 84 and 33% conditions showed extensive differences across a wide swath of the scalp in the control group. These onset at ~300 ms and extended to 750 ms, picking up again in the ~775 to 850 ms period, and showing a third voley of activity in the 900 to 950 ms period. The autism group showed much more spatially and temporally circumscribed condition effects, with differences centered around frontocentral regions in the 350 to 550 ms period (Fig. S1)

### Behavioral Results

Mean RT for the control and autism groups was 330 and 349 ms, respectively. For both groups, RTs were fastest for the highest cue validity condition, and slowest for the lowest. For the control group these RT differences scaled with cue validity, increasing by ~20 ms as cue validity decreased (with mean increases of 16, 27, and 16 ms from 100 to 84%, 84 to 67% and 67 to 33%, respectively). A similar pattern was seen in the autism group, except that RT barely changed between the 84 and 67% conditions (with increases of 20, 02, and 20 ms from 100 to 84%, 84 to 67% and 67 to 33%, respectively) (Fig. 5). To assess this statistically, we first performed a linear mixed effect model for RT with an interaction term between group and probability condition. The model revealed both a significant effect of condition (ß=-96.37, SE=4.23, p<0.01) and a group-by-condition interaction (ß=-34.43, SE=6.10, p<0.01) while showing no effect of group (ß=-6.43, SE=183.55, p=0.97) (Table 2). Follow-up mixed-model tests revealed that the main condition effect was driven by all pairwise comparisons of probability condition, whereas the group-by-condition interaction was due to the 100%-67%, 100%-33%, 84%-67%, 84%-33% condition pairs, reflecting smaller differences in mean RTs between these conditions in autism (See Table 2 and Table S5).

**FIGURE 5:**
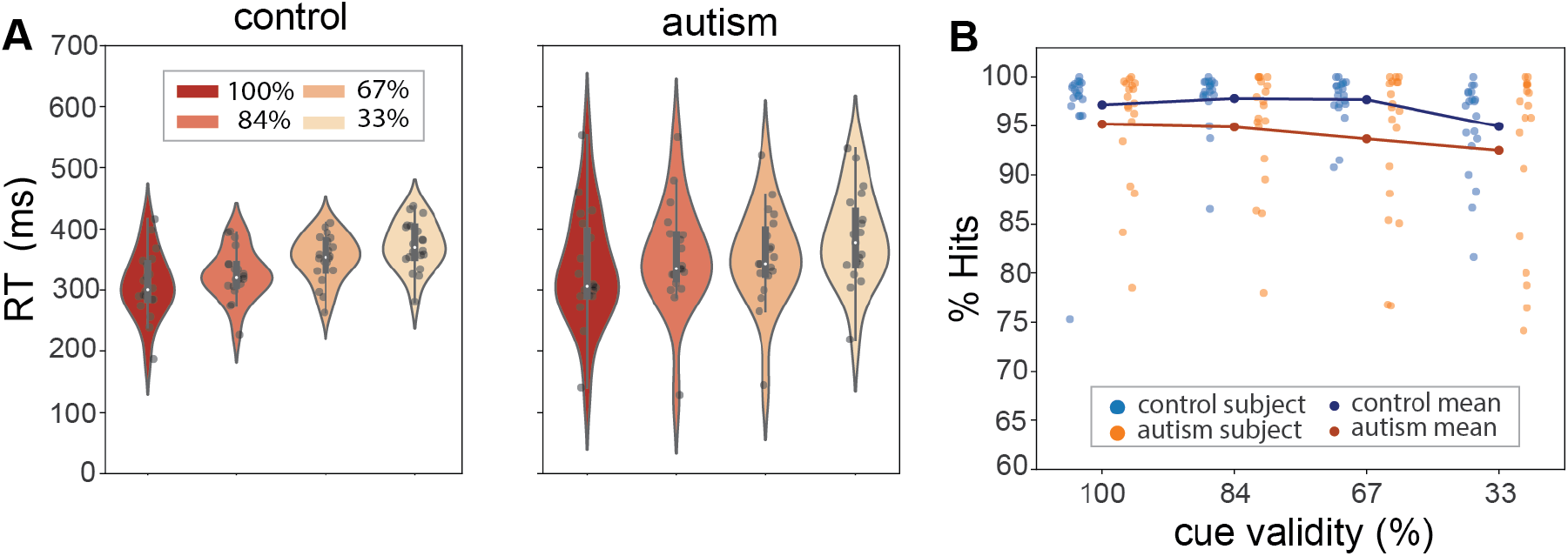
Reaction Time and Performance. (A) Target RTs in ms for the four probability conditions for control (left) and autism (bottom) groups. (B) Percent hit rate by probability condition. Dots that are connected by lines show averages. Each stand-alone dot represents an individual subject.

**TABLE 2:** Mixed Model Results for CNV, P3, SW, and RT. Group (Grp) = autism and neurotypical; Condition (Con) = probability condition; 100%, 84%, 67%, 33%).

|  | Coefficient | SE | z | P |
| --- | --- | --- | --- | --- |
| **CNV** |  |  |  |  |
| Intercept | -2.7 | 7.77 | -0.35 | 0.73 |
| Condition effect | 1.54 | 0.18 | 8.69 | <0.01 |
| Group effect | 1.38 | 10.99 | 0.13 | 0.9 |
| Con:Grp Interaction | -0.64 | 0.26 | -2.5 | 0.01 |
| **P3** |  |  |  |  |
| Intercept | 4.2 | 6.38 | 0.66 | 0.51 |
| Condition effect | -3.19 | 0.21 | -15.43 | <0.01 |
| Group effect | -0.43 | 9.02 | -0.05 | 0.96 |
| Con:Grp Interaction | 0.14 | 0.3 | 0.46 | 0.65 |
| **SW** |  |  |  |  |
| Intercept | 2.96 | 7.98 | 0.37 | 0.71 |
| Condition effect | -1.44 | 0.26 | -5.59 | <0.01 |
| Group effect | -0.86 | 11.29 | -0.08 | 0.94 |
| Con:Grp Interaction | 1.72 | 0.38 | 4.59 | <0.01 |
| **RT** |  |  |  |  |
| Intercept | 399.55 | 129.23 | 3.09 | <0.01 |
| Condition effect | -96.38 | 4.23 | -22.76 | <0.01 |
| Group effect | -6.43 | 183.55 | -0.03 | 0.97 |
| Con:Grp Interaction | 34.43 | 6.10 | 5.63 | <0.01 |

We examined the relationship between our neural and RT measures of flexibility in certainty of predictions (flexibility index: difference between 84% and 33% conditions) and SRS scores. These analyses were performed on a subset of the data due to missing SRS scores from 10 participants (5 each from the control and autism groups). We found significant correlations for the CNV (r(28) = 0.49, p < 0.01), P3 (r(28) = −0.31, p = 0.04), and SW (r(28) = −0.33, p = 0.03), whereas no significant correlation was found for RT (r(28) = −0.14, p = 0.22).

Both groups performed the task with high accuracy (96% and 93% respectively for control and autism groups; see Fig. 6). Mean hit rate to targets for the control group was more than 97% in the three highest cue validity conditions, and 94% for the lowest cue validity condition. For the autism group, hit rates decreased as cue validity decreased, from 95% to 92%. Statistical analyses revealed a main effect of condition effect (ß=0.02, SE<0.01, p<0.01) and a group by condition interaction (ß=0.02, SE<0.01, p=0.03; see Table S6).

## DISCUSSION

> *“For those of us who are on the spectrum, almost everything is black or white.”*
>
> — -Greta Thunberg (Thunberg, 2018)

We investigated how young adults with and without autism adjust prediction certainty, a central feature of predictive processing, upon parametric manipulation of cue validity ranging from 33% to 100%. Three distinct brain processes served to index the anticipation of temporally predictable targets (CNV), the evaluation and registration of target events (P3), and the updating of internal models (SW). Whereas the control group showed graded modulation of these brain responses and RT that was proportional to the level of cue validity, this pattern was not uniformly evident in the autism group. In particular, for the CNV (Fig. 2), there was a pronounced difference between the fully predictable condition (100% cue validity) and the less predictable conditions, whereas differences among the three less predictable conditions was substantially reduced. These CNV data suggest that autistic individuals do not modulate certainty of their predictions based on cue validity in the same highly flexible and reliable manner as do controls. Instead, the current data suggest that in autism certainty of a prediction is more binary (it’s either *certain* or *uncertain*) than graded, at least when faced with a changing and uncertain environment. Arguably, outsized responses to small deviations from what is expected (i.e., the 84% condition) could lead to the insistence on sameness phenotype, in which rules and routines are perpetually sought in everyday life, whereas the reduced differentiation among the 84, 67, and 33% cue validity conditions may relate to difficulty applying nuanced predictive information under ambiguous situations, such as those encountered in complex everyday social interactions.

The target P3, in contrast to the CNV, systematically modulated by cue validity not only for the control but also for the autism group (Fig. 3). This finding aligns with studies showing that autistic individuals represent stimulus statistics in a typical manner (Cannon et al., 2021; Knight et al., 2020; Manning et al., 2017), although it should be noted that condition effects were less robust in the autism group (see Fig. S1). The finding of relatively intact P3 modulation combined with impaired CNV modulation suggests that while stimulus statistics are calculated, the application of these stimulus statistics to prediction certainty is impaired. Future work is needed to determine if this finding is specific to environments where cue-target contingencies change relatively frequently, as in the present study, or if it represents a more generalized mode of operation in making predictions.

The CNV results appear to fit well with the theory of Highly Inflexible and Precise Prediction Errors in Autism (HIPPEA) proposed by Van de Cruys and colleagues (Van de Cruys et al., 2014). This theory posits that an atypically high level of precision is assigned to prediction errors in autism, by which even little variances in the environment will induce an update in the predictive model; this in turn leads to overfitted models, as even insignificant details/changes are considered important and reacted to, rather than being disregarded. Thus, with more precise prediction errors, even small changes evoke a large response, much as we see in the CNV responses for the autism group. Our data further suggest that prior empirical findings of reduced differentiation among different levels of cue validity (Lawson et al., 2017; Perrykkad et al., 2021) may not be indicative of reduced surprise, but rather of reduced flexibility in the representation of uncertainty. Whereas a 100% cue valid condition was not included in these studies, it clearly provides an important comparison when evaluating the magnitude of representation of uncertainty in predictions.

The behavioral data were also consistent with altered modulation of prediction certainty in autism. Whereas mean RT followed the expected pattern in the control group such that responses were faster when cue validity was higher and slower when it was lower (Fig. 5), mean RT differences between conditions were uniformly and statistically smaller in autism, and there was no RT difference at all between the intermediate conditions (84 and 67%). This attenuation of cue-validity effects on RT was present despite overall similar RTs for the autism and control groups (349 versus 330 ms).

Inspection of the ERPs revealed an additional response of interest, a positive-going distribution over frontal scalp 650-850 ms post target stimulus that, like the CNV and P3, varied in amplitude with respect to cue validity in the control group. This resembled the Slow Wave (SW) response that that has been highlighted in prediction tasks (de Gee et al., 2021; Loveless et al., 1987; Ruchkin et al., 1980), has a frontal-maximum topography (Loveless et al., 1987), and peaks between 500-800 ms after the event that follows a cue (de Gee et al., 2021; Sambrook et al., 2018). Even though the functional role of this component is debated, the observation that the SW is present during later stages of information processing has been taken to suggest that it may reflect an indepth analyses or re-evaluation process (Karniski et al., 1993), or a need for further processing (Ruchkin et al., 1980). In the context of the current study, the SW response may reflect participants’ re-evaluation and updating of the internal model of cue-target contingencies. In the control group, SW in response to targets was largest in amplitude for the 2 lowest cue validity conditions and smallest for the 100% condition (Fig. 4). This systematic pattern was not as evident in the autism group (Fig. 4), suggesting that autistic individuals do not update their internal model in a typical manner, after registering outcomes (Coll et al., 2020; Van de Cruys et al., 2013; Van de Cruys et al., 2017; Vishne et al., 2021). Figure 4B illustrates that numerically, for controls, SW increases systematically as target probability decreases, whereas in autism the difference was biggest between the 100% and the 84% conditions.

Of vital interest is whether and how these electrophysiological and behavioral indices of flexibility of predictions map onto the autism phenotype. To begin to address this question, we focused on SRS scores. The SRS scores provide a continuous measure of characteristics associated with the autism phenotype in the broader population as well as in autism (Constantino, 2013). These were significantly associated with reduced flexibility in representing prediction certainty (as measured by CNV, P3 and SW response differences between the 84% and 33% conditions). Although this requires replication in larger samples, these data provide preliminary evidence that impaired predictive processing may contribute to social difficulties and other behaviors associated with autism.

While our approach cannot identify the precise locus of disrupted processing, prior studies suggest several cortical/subcortical regions that contribute to CNV generation and the modulation of prediction certainty. For example, the anterior cingulate cortex (ACC) monitors the likelihood of events (Brown & Braver, 2005), is consistently highlighted in probabilistic tasks in functional imaging (Agam et al., 2010; O’Reilly et al., 2013) and animal studies (Stolyarova et al., 2019), and is thought to contribute to the CNV response (Gómez et al., 2003; Mulert et al., 2004; Nagai et al., 2004). The thalamus has also been implicated in the representation of precision in the context of predictive models (Kanai et al., 2015), and has been shown to contribute to trial-by-trial modulation of CNV amplitude (Nagai et al., 2004). Likewise, the prefrontal cortex is implicated in the representation of basic and more abstract prediction errors (Alexander & Brown, 2018; Zarr & Brown, 2016), and contributes to the CNV response (Gómez et al., 2007; Gómez et al., 2003; Mulert et al., 2004; Scheibe et al., 2010). Compellingly, activity in all of these brain regions has been shown to differ in autism (Balsters et al., 2016; Di Martino et al., 2009; Solomon et al., 2015; Tomasi & Volkow, 2019). Nevertheless, future studies using functional magnetic resonance imaging (fMRI) or intracranial EEG will be essential to identifying the network that underlies atypical representation of certainty in autism.

Finally, we should note that to understand whether there is a causal role between altered predictive processes and autism, it will be informative to assess at-risk populations (e.g., siblings of individuals diagnosed with autism) before the emergence of autism symptomatology, during infancy/early childhood (<2 years of age; e.g., see (Constantino et al., 2021)). For this, it will be necessary to design robust experimental assays of altered predictive processing for administration to very young children and lower functioning individuals. Understanding the exact problems with predictive processing is critical to the development of biomarkers for autism characteristics, and informing targeted therapies such as cognitive-behavioral approaches to helping affected individuals make more flexible predictions in everyday life.

## ACKNOWLEDGMENTS

We are grateful to the individuals who participated in this research and their families for their time and their commitment to the advancement of scientific discovery; without them this work would not be possible. We would like to thank Dr. Catherine Sancimino and Dr. Juliana Bates, who administered or supervised the clinical assessments, Dr. Ana Francisco for her suggestions on statistical analyses, Dr. Jose Luis Pena, Dr. Ruben Coen Cagli, and Dr. Eric Hollander for their valuable inputs at the student advisory committee meetings. We are also grateful to the research assistants and technicians at the Cognitive Neurophysiology lab of Albert Einstein College of Medicine who contributed to the collection of high-quality EEG data. The Human Clinical Phenotyping Core, where the majority of the children enrolled in this study were clinically evaluated, is a facility of the Rose F. Kennedy Intellectual and Developmental Disabilities Research Center (IDDRC) which is funded through a center grant from the Eunice Kennedy Shriver National Institute of Child Health & Human Development (NICHD U54 HD090260; P50 HD105352).

