## supplemental figures and tables for "All-or-None Evaluation of Prediction Certainty in Autism"

### SUPPLEMENTARY FIGURES

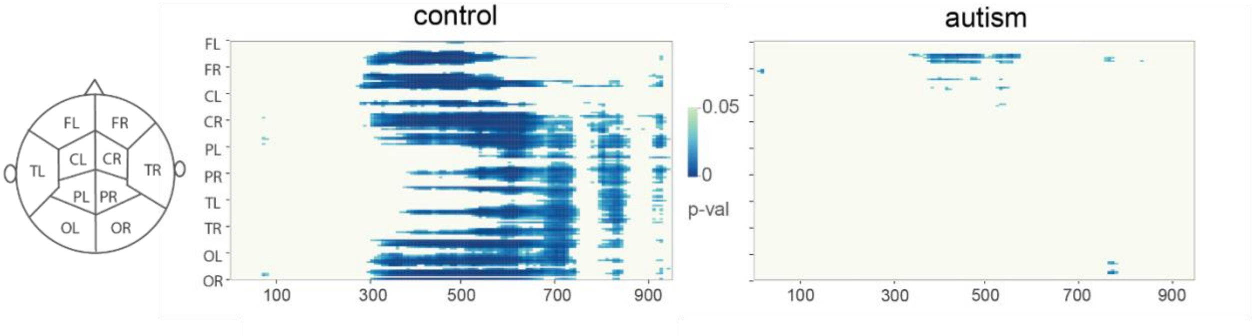


**SUPPLEMENTARY FIGURE 1: Statistical cluster plots constrasting target responses to 84% and 33% conditions.** The results of running t-tests across between 84% and 33% conditions over all channels and time points are displayed as an intensity plot. The x and y axes represent time and electrode location respectively. The color represents the p value for each data point.

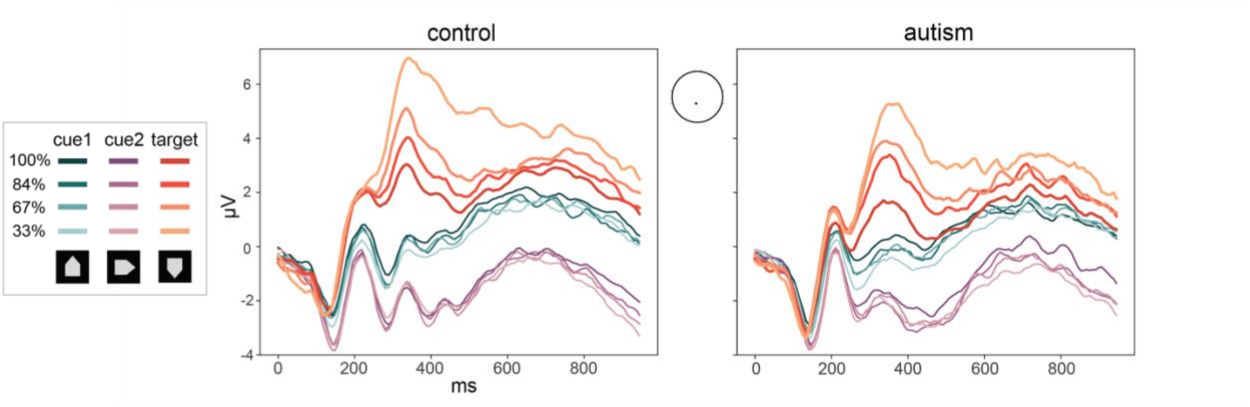


**SUPPLEMENTARY FIGURE 2: ERPs to cue1, cue2, and target.** ERP waveforms showing responses to cue1 (blue), cue2 (purple), and targets (red) at Cz for each of the probability conditions for control and autism participants.

### SUPPLEMENTARY TABLES

**SUPPLEMENTARY TABLE 1: Stimulus Statistics:** The number of each stimulus type is shown for a mini-block in each condition.

| Stimulus Type | 100% | 84% | 67% | 33% |
|---:|---:|---:|---:|---:|
| Item 1 | 24 | 24 | 24 | 24 |
| Item 2 | 24 | 24 | 24 | 24 |
| Target | 24 | 20 | 16 | 8 |
| Catch | 8 | 12 | 16 | 24 |
| Filler | 8 | 8 | 8 | 8 |
| Invalid Item | 0 | 4 | 8 | 16 |
| Total | 88 | 92 | 96 | 104 |

**SUPPLEMENTARY TABLE 2: Results of CNV Mixed Models for Pairwise Contrasts of Cue Validity Conditions.** Post-hoc mixed model results for each pairwise comparison of all cue validity conditions (100%-84%, 100%-67%, 100%-33%, 84%,67%, 84%-33%, 67%-33%).

| | Coefficient | SE | z | P |
|---|---|---|---|---|
| **100%-84%** | | | | |
| Intercept | -3.07 | 7.66 | -0.4 | 0.69 |
| Condition effect | 1.88 | 0.82 | 2.3 | 0.02 |
| Group effect | -0.07 | 10.83 | -0.01 | <0.01 |
| Con:Grp Interaction | 0.95 | 1.18 | 0.81 | 0.42 |
| **100%-67%** | | | | |
| Intercept | -1.97 | 7.48 | -0.26 | 0.79 |
| Condition effect | 0.78 | 0.36 | 2.15 | 0.03 |
| Group effect | 0.38 | 10.58 | 0.04 | 0.97 |
| Con:Grp Interaction | 0.5 | 0.53 | 0.94 | 0.35 |
| **100%-33%** | | | | |
| Intercept | -2.8 | 7.96 | -0.35 | 0.73 |
| Condition effect | 1.61 | 0.19 | 8.63 | <0.01 |
| Group effect | 1.5 | 11.26 | 0.13 | 0.89 |
| Con:Grp Interaction | -0.62 | 0.27 | -2.32 | 0.02 |
| **84%-67%** | | | | |
| Intercept | -1.33 | 7.51 | -0.18 | 0.86 |
| Condition effect | -0.19 | 0.78 | -0.25 | 0.8 |
| Group effect | 0.65 | 10.62 | 0.06 | 0.95 |
| Con:Grp Interaction | 0.1 | 1.13 | 0.09 | 0.93 |
| **84%-33%** | | | | |
| Intercept | -2.77 | 8.11 | -0.34 | 0.73 |
| Condition effect | 1.53 | 0.29 | 5.31 | <0.01 |
| Group effect | 1.66 | 11.47 | 0.15 | 0.89 |
| Con:Grp Interaction | -1.11 | 0.41 | -2.71 | 0.01 |
| **67%-33%** | | | | |
| Intercept | -3.08 | 7.94 | -0.39 | 0.7 |
| Condition effect | 2.47 | 0.42 | 5.86 | <0.01 |
| Group effect | 1.88 | 11.23 | 0.17 | 0.87 |
| Con:Grp Interaction | -1.77 | 0.61 | -2.92 | <0.01 |

**SUPPLEMENTARY TABLE 3: Results of P3 Mixed Models for Pairwise Contrasts of Cue Validity Conditions.** Post-hoc mixed model results for each pairwise comparison of all cue validity conditions (100%-84%, 100%-67%, 100%-33%, 84%,67%, 84%-33%, 67%-33%).

| | Coefficient | SE | z | P |
|---|---|---|---|---|
| **100%-84%** | | | | |
| Intercept | 3.97 | 6.38 | 0.62 | 0.53 |
| Condition effect | -2.94 | 0.73 | -4.02 | <0.01 |
| Group effect | 1.81 | 9.02 | 0.2 | 0.84 |
| Con:Grp Interaction | -2.23 | 1.06 | -2.11 | 0.04 |
| **100%-67%** | | | | |
| Intercept | 4.13 | 6.25 | 0.66 | 0.51 |
| Condition effect | -3.1 | 0.35 | -8.83 | <0.01 |
| Group effect | 0.26 | 8.84 | 0.03 | 0.98 |
| Con:Grp Interaction | -0.68 | 0.52 | -1.31 | 0.19 |
| **100%-33%** | | | | |
| Intercept | 4.23 | 6.84 | 0.62 | 0.54 |
| Condition effect | -3.21 | 0.24 | -13.51 | <0.01 |
| Group effect | -0.72 | 9.68 | -0.07 | 0.94 |
| Con:Grp Interaction | 0.3 | 0.34 | 0.87 | 0.39 |
| **84%-67%** | | | | |
| Intercept | 4.23 | 5.7 | 0.74 | 0.46 |
| Condition effect | -3.25 | 0.7 | -4.68 | <0.01 |
| Group effect | -0.65 | 8.07 | -0.08 | 0.94 |
| Con:Grp Interaction | 0.7 | 1.01 | 0.69 | 0.49 |
| **84%-33%** | | | | |
| Intercept | 4.26 | 6.63 | 0.64 | 0.52 |
| Condition effect | -3.29 | 0.33 | -9.86 | <0.01 |
| Group effect | -0.98 | 9.38 | -0.11 | 0.92 |
| Con:Grp Interaction | 1.09 | 0.48 | 2.27 | 0.02 |
| **67%-33%** | | | | |
| Intercept | 4.27 | 6.46 | 0.66 | 0.51 |
| Condition effect | -3.31 | 0.51 | -6.47 | <0.01 |
| Group effect | -1.05 | 9.14 | -0.12 | 0.91 |
| Con:Grp Interaction | 1.31 | 0.75 | 1.75 | 0.08 |

**SUPPLEMENTARY TABLE 4: Results of SW Mixed Models for Pairwise Contrasts of Cue Validity Conditions.** Post-hoc mixed model results for each pairwise comparison of all cue validity conditions (100%-84%, 100%-67%, 100%-33%, 84%,67%, 84%-33%, 67%-33%).

| | Coefficient | SE | z | P |
|---|---|---|---|---|
| **100%-84%** | | | | |
| Intercept | 3.53 | 7.84 | 0.45 | 0.65 |
| Condition effect | -2.09 | 0.9 | -2.32 | 0.02 |
| Group effect | 1.86 | 11.09 | 0.17 | 0.87 |
| Con:Grp Interaction | -1.06 | 1.3 | -0.82 | 0.41 |
| **100%-67%** | | | | |
| Intercept | 3.99 | 7.93 | 0.5 | 0.62 |
| Condition effect | -2.55 | 0.45 | -5.72 | <0.01 |
| Group effect | -1.94 | 11.21 | -0.17 | 0.86 |
| Con:Grp Interaction | 2.74 | 0.66 | 4.19 | <0.01 |
| **100%-33%** | | | | |
| Intercept | 2.59 | 8.26 | 0.31 | 0.75 |
| Condition effect | -1.15 | 0.29 | -4.03 | <0.01 |
| Group effect | -0.59 | 11.68 | -0.05 | 0.96 |
| Con:Grp Interaction | 1.38 | 0.42 | 3.34 | <0.01 |
| **84%-67%** | | | | |
| Intercept | 4.26 | 7.61 | 0.56 | 0.58 |
| Condition effect | -2.96 | 0.93 | -3.2 | <0.01 |
| Group effect | -4.18 | 10.76 | -0.39 | 0.7 |
| Con:Grp Interaction | 6.12 | 1.35 | 4.53 | <0.01 |
| **84%-33%** | | | | |
| Intercept | 2.5 | 8.09 | 0.31 | 0.76 |
| Condition effect | -0.86 | 0.41 | -2.12 | 0.03 |
| Group effect | -0.84 | 11.44 | -0.07 | 0.94 |
| Con:Grp Interaction | 2.15 | 0.59 | 3.66 | <0.01 |
| **67%-33%** | | | | |
| Intercept | 2.12 | 8.36 | 0.25 | 0.8 |
| Condition effect | 0.29 | 0.66 | 0.43 | 0.67 |
| Group effect | -0.12 | 11.83 | -0.01 | 0.99 |
| Con:Grp Interaction | -0.02 | 0.97 | -0.02 | 0.99 |

**SUPPLEMENTARY TABLE 5: Results of Reaction Time Mixed Models for Pairwise Contrasts of Cue Validity Conditions.** Post-hoc mixed model results for each pairwise comparison of all cue validity conditions (100%-84%, 100%-67%, 100%-33%, 84%,67%, 84%-33%, 67%-33%).

| | Coefficient | SE | z | P |
|---|---|---|---|---|
| **100%-84%** | | | | |
| Intercept | 399.38 | 136.09 | 2.93 | <0.01 |
| Condition effect | -97.57 | 15.66 | -6.23 | <0.01 |
| Group effect | 54.4 | 192.49 | 0.28 | 0.77 |
| Con:Grp Interaction | -27.59 | 22.52 | -1.22 | 0.22 |
| **100%-67%** | | | | |
| Intercept | 426.89 | 134.91 | 3.16 | <0.01 |
| Condition effect | -125.09 | 7.6 | -16.45 | <0.01 |
| Group effect | -33.4 | 190.85 | -0.17 | 0.86 |
| Con:Grp Interaction | 60.21 | 11.11 | 5.41 | <0.01 |
| **100%-33%** | | | | |
| Intercept | 390.13 | 135.82 | 2.87 | <0.01 |
| Condition effect | -88.32 | 4.75 | -18.59 | <0.01 |
| Group effect | 0.98 | 192.19 | <0.01 | 0.99 |
| Con:Grp Interaction | 25.82 | 6.83 | 3.78 | <0.01 |
| **84%-67%** | | | | |
| Intercept | 443.04 | 121.47 | 3.64 | <0.01 |
| Condition effect | -149.55 | 14.84 | -10.07 | <0.01 |
| Group effect | -84.92 | 171.83 | -0.49 | 0.62 |
| Con:Grp Interaction | 138.27 | 21.55 | 6.41 | <0.01 |
| **84%-33%** | | | | |
| Intercept | 389.17 | 119.55 | 3.25 | <0.01 |
| Condition effect | -85.42 | 6.06 | -14.08 | <0.01 |
| Group effect | -4.54 | 169.08 | -0.02 | 0.97 |
| Con:Grp Interaction | 42.58 | 8.7 | 4.89 | <0.01 |
| **67%-33%** | | | | |
| Intercept | 377.63 | 116.6 | 3.23 | <0.01 |
| Condition effect | -50.45 | 9.3 | -5.42 | <0.01 |
| Group effect | 12.67 | 164.9 | 0.07 | 0.93 |
| Con:Grp Interaction | -9.6 | 13.47 | -0.71 | 0.47 |

**SUPPLEMENTARY TABLE 6: Mixed Models Results for Percent Hits.** Mixed model results for comparisons of cue validity conditions.

| | Coefficient | SE | z | P |
|---|---|---|---|---|
| **100%-84%-67%-33%** | | | | |
| Intercept | 0.95 | 0.20 | 4.7 | <0.01 |
| Condition effect | 0.02 | <0.01 | 3.74 | <0.01 |
| Group effect | -0.04 | 0.28 | -0.14 | 0.88 |
| Con:Grp Interaction | 0.02 | <0.01 | 2.16 | 0.03 |
| **100%-84%** | | | | |
| Intercept | 1.01 | 0.19 | 5.25 | <0.01 |
| Condition effect | -0.04 | 0.02 | -2.05 | 0.04 |
| Group effect | -0.08 | 0.27 | -0.29 | 0.76 |
| Con:Grp Interaction | 0.06 | 0.03 | 2.00 | 0.04 |
| **100%-67%** | | | | |
| Intercept | 0.98 | 0.19 | 4.97 | <0.01 |
| Condition effect | -0.01 | 0.01 | -1.16 | 0.24 |
| Group effect | -0.08 | 0.27 | -0.28 | 0.77 |
| Con:Grp Interaction | 0.06 | 0.01 | 3.82 | <0.01 |
| **100%-33%** | | | | |
| Intercept | 0.93 | 0.20 | 4.55 | <0.01 |
| Condition effect | 0.03 | <0.01 | 4.65 | <0.01 |
| Group effect | -0.02 | 0.29 | -0.09 | 0.92 |
| Con:Grp Interaction | 0.01 | 0.01 | 0.98 | 0.32 |
| **84%-67%** | | | | |
| Intercept | 0.96 | 0.19 | 4.88 | <0.01 |
| Condition effect | 0.01 | 0.02 | 0.67 | 0.5 |
| Group effect | -0.07 | 0.27 | -0.28 | 0.77 |
| Con:Grp Interaction | 0.05 | 0.03 | 1.72 | 0.08 |
| **84%-33%** | | | | |
| Intercept | 0.92 | 0.21 | 4.41 | <0.01 |
| Condition effect | 0.05 | 0.01 | 5.47 | <0.01 |
| Group effect | -0.02 | 0.29 | -0.07 | 0.93 |
| Con:Grp Interaction | <0.01 | 0.01 | -0.45 | 0.64 |
| **67%-33%** | | | | |
| Intercept | 0.92 | 0.22 | 4.16 | <0.01 |
| Condition effect | 0.08 | 0.01 | 4.6 | <0.01 |
| Group effect | -0.01 | 0.31 | -0.03 | 0.97 |
| Con:Grp Interaction | -0.04 | 0.02 | -1.71 | 0.08 |
